# Three complete genome sequences of Penguin megrivirus from ornithogenic soil, Adelie penguin and Weddell seal of Antarctica

**DOI:** 10.1101/858043

**Authors:** Ashutosh Aasdev, Anamika Mishra, Manoj Nair, Satyam D Pawar, Chandan K Dubey, Sandeep Bhatia, Prabir G Dastidar, Vijendra Pal Singh, Ashwin Ashok Raut

**Author notes:** Corresponding Author: Ashwin Ashok Raut,.

## Abstract

We report the complete genome sequences of Penguin megriviruses from three different sources from Antarctica namely feces of Adelie penguin, feces of Weddell seal and ornithogenic soil. Phylogenetic analysis indicates the prevalence of very similar viruses in different sources of Antarctic environment. These genome sequences aid to understand the evolution of megriviruses in Antarctic ecology and reveal their place in global megrivirus phylogeny.

## Announcement

The genus *Megrivirus* belongs to positive sense single stranded RNA virus family *Picornaviridae*. Megrivirus was classified as a genus in 2013 (1). Previously the genus had only one species, the Turkey hepatitis virus, but identification of new viruses has expanded the genus to five species and many unclassified viruses. All megriviruses identified infect aves, and have been identified from chicken to wild birds (2). Penguin megrivirus belongs to species Megrivirus E and was only recently reported in 2017 (3).

Samples for this study were collected in February 2018 as part of the 37^th^ Indian Scientific Expedition to Antarctica, Ministry of Earth Sciences, Government of India. RNA extraction from the fecal samples was done on site at Bharati Station, Antarctica. Fresh fecal samples of Adelie penguin and Weddell seal were suspended in PBS and centrifuged at 12,000 RCF for 15 min. The supernatant was filtered (0.45µm) and RNA was extracted using QIAamp Viral RNA mini kit without adding carrier RNA. The RNA samples were dried in GenTegra RNA plates and transported to NIHSAD, Bhopal, India at room temperature. Ornithogenic soil sample transported in cold chain, was processed in the BSL3+ containment lab at NIHSAD. One gram of soil was suspended in 10 mL HBSS and incubated at room temperature for an hour followed by centrifugation at 6,000 RCF for 15 minutes at 4°C. The supernatant was filtered through 0.45μm filter and ultracentrifuged at 180,000 RCF for 4 hours at 4°C. The pellet was resuspended in 280 μL HBSS and viral RNA extraction was done using QIAamp Viral RNA Mini Kit.

Highthroughput sequence data was generated at Genotypic Technology Private Limited, Bangalore, India by 2 × 150 bp paired end Illumina sequencing on MiSeq platform, using NEBNext^®^ Ultra™ Directional RNA Library Prep Kit. The raw reads were quality controlled using FastQC v0.11.3 (https://www.bioinformatics.babraham.ac.uk/projects/fastqc/). Adapter and low quality bases were removed using Trimmomatic v0.39 (4). BLASTn of the processed reads was performed against a custom viral genome database using local blast v2.8.1+. Reads for individual virus reference sequences were extracted. The reference sequences with high number of reads originating from them were used for mapping the reads using BWA-MEM v0.7.17-r1194-dirty (5). Consensus sequence were evaluated using BCF tools v1.9 (6). PCR confirmation of virus genome was done using primers, PM-F120 5’-TGGAGCTGCCATCACGTGTT-3’ and PM-R120 5’-TCCTGACACTGGTCACGTCT-3’. Few of the terminal nucleotides could not be read for the sequences as indicated in Table 1. NCBI ORF finder was used to predict the ORF’s. The polyprotein ORF is between nucleotide positions 718 and 9030 and codes for a 2770 amino acid polypeptide. The complete polypeptide sequences of 16 megriviruses was downloaded from NCBI and protein alignment performed using MUSCLE in MEGA7 (7). Neighbour-joining phylogenetic tree was constructed using Poisson model and 1000 bootstrap replicates. The viruses form a monophyletic group. The pairwise amino acid identity amongst the viruses under study is from 99.75% to 99.9%. The 5’ UTR secondary structure was predicted using mfold (8). The structure contains elements similar to domain II and III of megrivirus IRES (9, 10). Also, the conserved 20 nucleotides which make the ‘8’ like structure in domain III are present (9, 10). The 3’ UTR contains 2 unit A repeats while the AUG rich region following the repeats is absent (10).

**Table 1:** Sampling location and sequencing metrics of the Penguin megriviruses from Antarctica.

| **Sample** | **Accession Numbers** | **Location (Coordinates)** | **Total number of reads** | **Number of mapped reads** | **Average coverage** | **Percentage genome obtained** | **Number of missing terminal nucleotides** | **G+C content (%)** |
| --- | --- | --- | --- | --- | --- | --- | --- | --- |
| Ornithogenic soil | MN453780 | Svenner Islands, Prydz Bay (-69.03, 76.83) | 176,926,262 | 4,273,385 | 62,167 | 99.93 | 4 | 42.87 |
| Adelie penguin faeces | MN453781 | Svenner Islands, Prydz Bay (-69.03, 76.83) | 144,867,534 | 11,520 | 169 | 99.88 | 13 | 43.03 |
| Weddell seal faeces | MN453782 | Larsemann Hills, Prydz Bay (-69.58, 78.55) | 133,147,552 | 3,085 | 44 | 99.92 | 8 | 43.04 |

In summary we report the complete genome sequences and phylogenetic relation of closely related Penguin megiviruses from three different sources of Antarctica samples. The presence of the virus in fecal samples points to its probable enteric localization. Also, Penguin megrivirus, which is known to infect birds, was identified in Weddell seal feces and there is further need to investigate its ecological relations and infectivity.

## Data Availability

The genome nucleotide sequences have been submitted to GenBank under accession numbers MN453780, MN453781 and MN453782.

## Acknowledgements

Indian Scientific Expeditions to Antarctica (ISEA) by National Center for Polar and Ocean Research, Goa suported by Earth System Science Organization, Ministry of Earth Sciences (ESSO-MoES), Government of India (GoI). Indian Council of Agricultural Research and Director, National Institute of High Security Animal Diseases, Bhopal for deputation of Scientist under ISEA and providing necessary facilities to conduct this work.

